# Host-enemy interactions provide limited biotic resistance for a range-expanding species via reduced apparent competition

**DOI:** 10.1101/2022.06.28.498037

**Authors:** Kirsten M. Prior, Dylan G. Jones, Shannon A. Meadley-Dunphy, Susan Lee, Alyson K. Milks, Sage Daughton, Andrew A. Forbes, Thomas H. Q. Powell

## Abstract

As species ranges shift in response to anthropogenic change, they lose coevolved or coadapted interactions and gain novel ones in recipient communities. Range-expanding species may lose or experience weak antagonistic interactions with competitors and enemies, and traits of interacting species will determine the strength of interactions. We leveraged a poleward range expansion of an oak gall wasp that co-occurs on its host plant with other gall wasp species and interacts with shared natural enemies (largely parasitoid wasps). We created quantitative host-parasitoid interaction networks by sampling galls on 400 trees. We compared network structure and function and traits of hosts and parasitoids in the native and expanded range. Interaction networks were less diverse in the expanded range, with low complementarity of parasitoid assemblages among hosts. While whole networks were more generalized in the expanded range, interactions with the range-expanding species were more specialized. This was not due to a loss of specialist enemies but weak apparent competition by shared generalist enemies. Phenological divergence of enemy assemblages attacking the novel and co-occurring hosts was greater in the expanded range that may contribute to weak apparent competition. Given the rate and extent of anthropogenic-driven range expansions, it is pressing to uncover how complex biotic interactions are reassembled.

## Introduction

Human activity is causing the reorganization of Earth’s biota as species are transported around the globe via trade and traffic and shift their ranges in response to climate and land-use change (1–3). When species move into new locations, interacting species are not likely to move in concert due to differences in dispersal or niche requirements (4–7). As a result, coevolved or coadapted interactions are lost, and novel associations are formed in new locations. This biogeographic flux disrupts complex networks of biotic interactions with cascading effects in ecosystems (6–9). Here, we uncover how changes in biotic interactions in networks of interacting species contribute to the dynamics of species’ range expansions, which is pressing given the extent and pace of anthropogenic change.

Changes in biotic interactions between species’ native and expanded ranges affect the population dynamics of species as they move into new regions. If antagonistic interactions with predators or competitors are lost or reduced, range-expanding species experience “high niche opportunities” (*sensu* (8); i.e., reduced competition leading to increased resources or reduced predation) that may lead to demographic release or increased fitness or population growth “ecological release” (6,8,10,11). Ultimately, net changes between the native and expanded range in biotic interactions (including indirect interactions) in the context of the abiotic environment will determine if range-expanding species experience high niche opportunities and ecological release. If net changes in interactions provide greater population control of range-expanders, they experience “biotic resistance,” with lower fitness or population growth in the expanded range (6,8,11).

The above-described community ecology framework is an explanation for why some introduced species become invasive (8,10,12–14) and it is more recently applied to species undergoing shorter-distance range expansions, including in response to climate change (6,7,15,16). It is predicted that differences in biotic interactions will be more significant and outcomes of those altered interactions more severe when species are moved over long distances (i.e., inter-continental introductions) into communities with which they share little or no coevolutionary history (11,17). Yet, there are growing examples of short-distance expanders (i.e., intra-continental expanders) experiencing ecological release (3,17–19). Short-distance expanders may experience release not just from coevolved or coadapted species, but also from coadapted populations (20–22), and especially when moving poleward may encounter a reduction in community diversity in northern locations (9,18,21,23). Moreover, instances of short-distance expansions are particularly tractable for testing hypotheses about the consequences of altered biotic interactions because the similarity of ecosystems and underlying species interaction networks over short distances allows for more direct comparisons of the biotic interactions affecting the focal species. Here, we leverage a short-distance poleward expansion of a phytophagous insect in a tractable host-enemy community to examine the community dynamics of short-distance expansions.

Parasitoid wasps are primary enemies of phytophagous insects and often interact in antagonistic networks (24–28). Species undergoing poleward range expansions may encounter less diverse communities in higher latitudes with weaker antagonistic interactions (29–31). If networks are also less specialized in poleward locations (32,33), recipient communities may provide limited biotic resistance as specialized antagonistic networks with high trophic complementarity are predicted to have high “function” or host control (34–36). Range-expanding insects may not only infiltrate less diverse or specialized recipient networks but also lose ancestral specialist enemies that fail to shift or lag behind range-expanding hosts (“enemy release”). Range-expanding species may also escape generalist enemies if generalists fail to follow, if fewer are in the recipient species pool, or if they fail to effectively switch from co-occurring hosts to the novel host (“release from apparent competition”) (18,21,37–39). Previous studies have revealed that range-expanding or introduced insects both lose specialist enemies from their native range, and even if generalist enemies attack novel hosts, that apparent competition is often weak due to reduced effectiveness on novel hosts (18,21,37–40).

Trait variation in one trophic level influences community assembly in interacting trophic levels (41,42). A lack of biotic resistance may result from recipient communities possessing divergent traits from range-expanding species that prevent effective host switching or sharing by enemies (42). Selection favors the evolution of defensive traits that reduce the success of parasitoid attacks in insect hosts. At the same time selection favors traits in parasitoids that help them evade host defenses (42–45). Traits include morphological features such as body size that can facilitate host defense or ovipositor size that can facilitate parasitoid attack (44–46). Phenology is also essential to interactions between insect hosts and parasitoids, as successful development for parasitoids requires that they attack hosts during discrete time windows, outside of which their development will not correspond with host resources (43,47). High niche opportunities and ecological release may occur when morphology and phenology of range-expanding hosts diverge from interacting host and enemy species in recipient communities.

A community of oak gall wasps (Hymenoptera: Cynipidae: Cynipini) co-occur on a dominant oak, *Quercus garryana*, in North American western oak ecosystems and are attacked by a community of natural enemies. One oak gall wasp species, *Neuroterus saltatorius*, hereafter “Nsal,” is expanding its range poleward, occurring at higher abundances in its expanded range, where it causes damage to *Q. garryana* (18,48,49). Oak gall wasps induce structures (galls) on plant tissue. Galls vary in traits, including size, shape, and texture, with several traits thought to be defensive adaptations to evade attack from natural enemies (43,50–52). Host gall morphology and phenology influence the assemblages of enemies attacking host species (42,51,53). Oak gall wasp-enemy communities are tractable multi-trophic communities that are excellent systems for uncovering direct and indirect trophic interactions and how interactions are altered under anthropogenic change (18,23,38,50–52,54,55).

To reveal how direct and indirect interactions among co-occurring hosts and natural enemies contribute to biotic resistance under poleward range expansions, we performed systematic surveys of oak gall wasps co-occurring on *Q. garryana* and their interacting natural enemies in the native and expanded range of Nsal. i) We created quantitative oak gall wasp-enemy interaction networks to compare differences in network structure (diversity and distribution) that relate to function (biotic resistance) between regions. ii) We calculated species-level metrics to uncover potential mechanisms of enemy loss under range expansions, that is, if Nsal loses interactions with putative specialist enemies or shared generalist enemies that attack co-occurring hosts. iii) We compared morphological traits and phenology of hosts and interacting enemies to uncover if trait divergence provides niche opportunities for the range-expanding host. iv) Finally, we measured the function of the host-enemy community, that we defined as the ability of the community to provide biotic resistance to the novel host. We predict that i) Nsal interacts with less diverse or specialized networks in the expanded range; ii) Nsal loses interactions with enemies from the native range that fail to follow or in the expanded range that fail to attack the novel host effectively; iii) weak biotic resistance is mediated by divergence in morphological traits and phenology between recipient host-enemy communities and the novel host; and iv) that interactions confer weaker biotic resistance (i.e., lower rates of successful enemy attack) in the expanded range. Uncovering how complex networks of biotic interactions are altered under anthropogenic change is essential (56,57) given the extent and pace of species’ range changes under anthropogenic change.

## Methods

### Study system

*Quercus garryana* Douglas ex. Hook (Fagaceae) ranges from northern California to Vancouver Island, British Columbia (BC) and is the only oak species from Oregon northwards. In western oak ecosystems, where no other oaks occur, *Q. garryana* is the predominant overstory woody vegetation. *Q. garryana*-ecosystems occur in the rain shadow of the coastal mountain ranges as savannas, grasslands, deep soil woodlands, or on rocky outcrops. *Q. garryana*-ecosystems become patchier at higher latitudes in northern Washington and Vancouver Island, BC (58).

Oak gall wasps (Hymenoptera: Cynipidae: Cynipini) are a specialized group of phytophagous insects that deposit their eggs in plant tissue of Fagaceae (oaks), inducing the formation of galls. Galls house and provide nutritive tissue to the gall wasp larvae during development (43,52). The majority of oak gall wasps have two generations, a gamic (sexual) and an agamic (asexual) generation that each form distinct galls (59). Gall structures vary among species and between generations and occur on various plant tissues (43,51,60). There are approximately 1000 oak gall wasp species, with the Nearctic having ∼700 species (60–62). Oak gall wasps support a rich community of natural enemies, predominantly parasitoid wasps in the superfamily Chalcidoidea. These wasps are often solitary ectoparasites that attack one to a few hosts (specialists) to multiple hosts (generalists), and largely only attack oak gall wasps or parasitoids attacking gall wasps (24,63,64). Parasitoid wasps that emerge out of galls are parasitoids that directly attack gall wasps or inquilines (other organisms that live inside galls or gall tissue) or are hyperparasitoids of parasitoids attacking gall wasps or inquilines (64).

*Neuroterus saltatorius* (Edwards) (*hereafter*, “Nsal”) induces galls on white oaks from Texas throughout the western portion of the United States, including *Q. garryana* (60,65). The native range of Nsal is restricted to mainland North America; however, in the early 1980s it expanded its range onto Vancouver Island, BC (48,49), which also defines the northernmost range of *Q. garryana* (18,55). Nsal’s early-spring gamic generation is a clustered integral leaf gall and its agamic generation occurs in the summer and is a detachable leaf gall. Both generations are approximately 1-2 mm in size (48). The detachable galls drop from the leaves in mid-late summer where they remain in the leaf litter for the winter. Adult gall wasps emerge the following spring (48). The agamic generation of Nsal occurs at higher abundance on *Q. garryana* in its expanded range, with higher frequency of infested trees. Some trees are infested in the native range of Nsal, but at low frequency (18,23,65). Particularly high abundances can cause foliar scorching with negative effects on oaks and oak associated species (18,48,49,55).

### Oak gall wasp and parasitoid enemy surveys

In 2017, we chose 10 study sites that were patches of oaks. We chose four sites in Nsal’s native range that were the largest oak patches with Nsal closest to its expanded range (18,23) and six sites in the expanded range that are some of the largest intact *Q. garryana* sites in the expanded range (Fig. 1). Sites were open oak grasslands or savannas with *Q. garryana* as the dominant tree, ranging in size from 6-130 ha and separated by at least 10 km in a matrix of rural agriculture, residential areas, and *Pseudotsuga menziesii* forests (Fig. 1, S1_Table S1).

**Figure 1:**
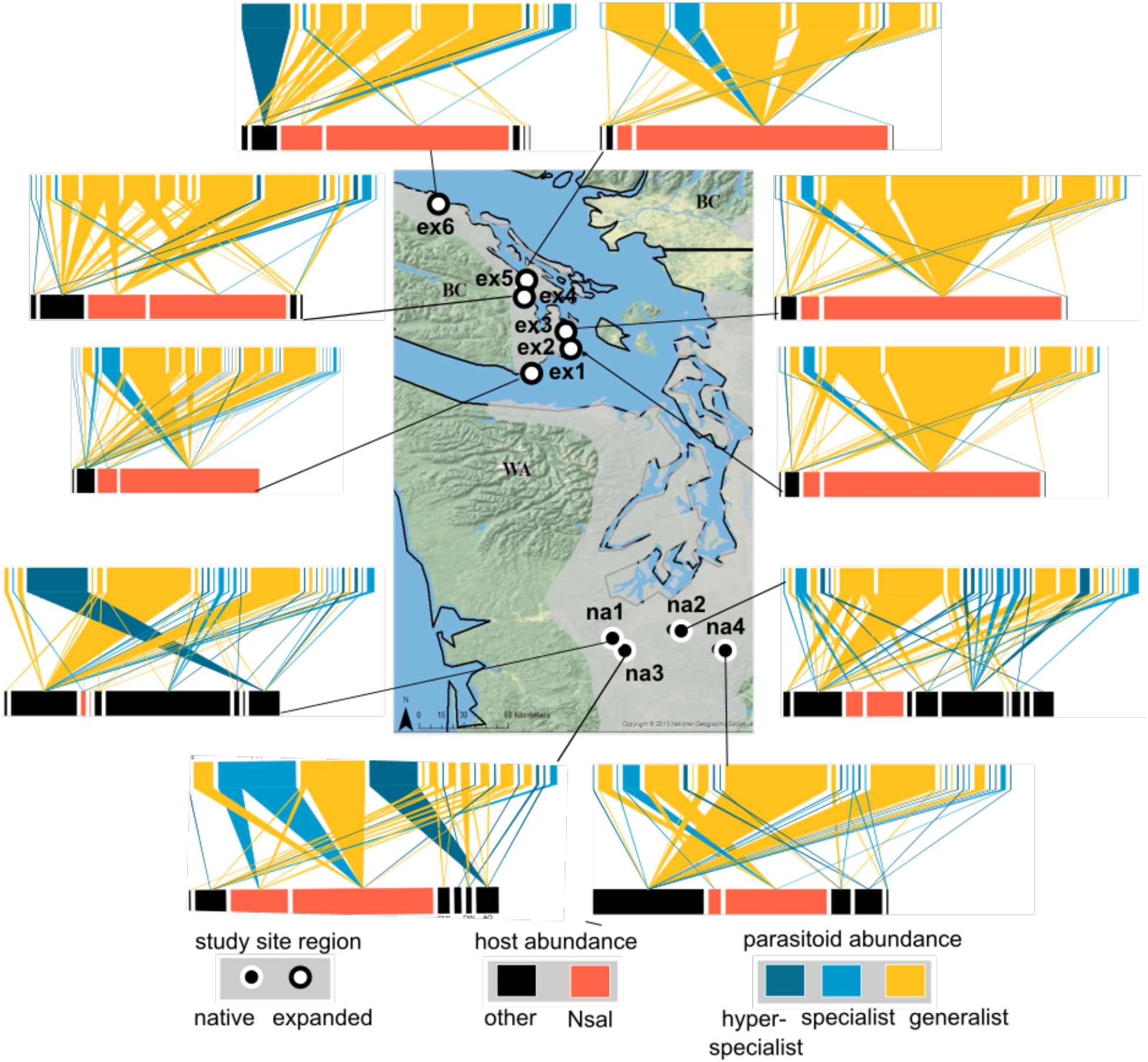
Range of *Quercus garryana* (shaded gray) in northern Washington State and southern Vancouver Island, British Columbia. At each study site in the native range of Nsal (dark symbols) and expanded range (white symbols) developed galls were collected on 40 trees over four survey dates that span development time of Nsal. The bipartite quantitative networks depict interactions (links) between hosts (bottom bar) and emerged parasitoids (top bar). Blocks represent species and the width of the blocks reflect the relative abundance of hosts and of parasitoids emerged from each host. Networks were combined over survey periods. Native hosts to both regions depicted by black bars, and the range-expanding host by orange bars. Parasitoid types are depicted by colors: dark blue are hyperspecialists (attack 1 host), medium blue specialists (<3 hosts), or yellow generalists) (S1_Tables S2,S3).

At each study site, we performed surveys of oak gall wasps on *Q. garryana* during four separate sampling periods. The timing of surveys coincided with the two generations (gamic and agamic) of Nsal (mid-May to late-July). Surveys were conducted on a rotating basis every 10-12 days, starting with sites ex1-3, then ex4-6, then na1-4 (Fig. 1; S1_Table S1). This order was chosen because oaks in ex1-2 have earlier phenology than ex4-6 and na1-4 (18).

We surveyed 10 trees during each period, 40 trees per site. Trees were chosen haphazardly at a site that were at least 10 m apart, spreading out sampled trees among surveys. For a tree to be sampled, it had to be larger than 2 m in height, and we needed to have observed a gall wasp species within 5 minutes of searching reachable branches around the tree, using a 5 ft. ladder (up to ∼10 ft.). On 10 branches spread around the tree, we searched 10 leaf clusters for leaf galls and 1 m of branches for stem galls. All oak gall wasp individuals were identified via gall morphology, contacting experts in some cases (60,66) (S1_Table S2). We were able to identify gall morphotypes to species, except for *Disholcaspis mamillana* and *D. simulata*, and we lumped these two species together.

Mature galls were collected and stored in rearing containers, separated by gall morphotype, site, and survey date. For close to one year, galls were kept in environmental chambers set to summer Pacific northwest conditions (25° C, 14:10). These standard conditions were chosen because optimal environmental rearing conditions for each morphotype are unknown. Since parasitoids emerged out of all gall morphotypes, except for those in which only a small number of individuals were collected, was evidence that conditions were largely suitable. Once a week, containers were checked for emergents, which were collected and stored at -80 °C. We identified emergent wasps to family level first. Then for wasp families with known parasitoids, we identified individuals to the lowest taxonomic unit using taxonomic keys (67) and with the help of experts in some cases (18). We identified 63 unique parasitoid wasp morphospecies in 11 families in Superfamilies Chalcidoidea, Ichneumonoidea, and Platygastroidea (S1_Table S3). Inquiline wasps (i.e., cynipids that are not parasitoids, but feed on gall or plant tissue) also emerged from galls, but we did not include these in our networks as they may not all act as enemies (see S1_Supplementary Methods). While we included morphospecies in families with known parasitoid wasps, not all wasps may reflect a direct interaction with the gall former host. For example, they could be parasitoids of inquilines or hyperparasitoids (see S1 for details). Given that many direct associations of parasitoids are undescribed, we were conservative and included all morphospecies in our analysis from families with known parasitoids, assuming that most caused deaths (be it direct or indirect) of gall wasp hosts. Over 99% of parasitoid individuals reared were from taxonomic groups that are known to directly associate with gall wasps (see S1) (64,67).

### Host-parasitoid interaction networks

We created quantitative bipartite interaction networks consisting of bars (gall wasp host morphotypes and parasitoid morphospecies) and links (interactions) using the bipartite package in R (68). The width of the bottom bars represents the relative abundance of collected host morphotypes (Fig. 1, S2_Fig. S1). For multilocular galls (that contain multiple host individuals) (*N. washingtonensis, A. quercuscalifornicus*) (S1_Table S2), we multiplied each gall by an estimated number of wasp larvae in galls. We treated separate generations of Nsal as their own morphotypes, given that they occur at different times and may have different parasitoid assemblages, as is also found in other oak gall wasps (42). The width of the top bars and links represent relative parasitoid emergence frequency of each parasitoid morphospecies from each host morphotype.

We created bipartite quantitative interaction networks for each study site by pooling interactions among survey periods (Fig. 1). We created site-level rather than survey-level networks, given that survey-level networks are not independent. Several host species (including Nsal) occur throughout the four survey periods, and parasitoid interactions are linked over time, given that many parasitoids are likely multivoltine (63). We created regional networks (pooling sites within regions), and a metanetwork (pooling all sites) to perform network and trait analyses (S2_Fig. S1).

We calculated network-level metrics to describe differences in network structure and function between regions (S1_Table S4). We chose metrics that describe network diversity and distribution (i.e., specialization) that relate to uncovering the potential for host-parasitoid communities to provide biotic resistance. For metrics where weighting was possible, we weighted by interaction frequencies as weighted metrics represent functional importance of species and their interactions in networks and are more robust to sampling biases (69,70).

To compare network size and diversity, we estimated host morphotype and parasitoid morphospecies richness by calculating abundance-based Chao 1 estimates using the estimate function with the vegan package in R (71). We also estimated interaction richness as Chao 1 (as the number of unique interactions between hosts and parasitoids) (72) (see Supplementary Methods_S3). We calculated network size as the estimated number of hosts x parasitoids and included this as a factor in models (see below). We also calculated interaction diversity (Shannon Entropy, H_2_) and interaction evenness using weighted interaction diversity across networks (56).

To examine network distribution (i.e., specialization), we calculated the proportion of specialist parasitoid morphospecies (defined as parasitoids attacking three or fewer hosts in the metanetwork; (42) ; S2_Fig. S1) out of all parasitoid morphospecies in each site network. We also calculated network-level specialization H_2_^’^ that represents weighted network specialization or the exclusiveness of host-parasitoid interactions relative to each other (73). Weighted connectance was calculated as the frequency of realized interactions out of potential interactions or the linkage density (the number of interactions per species weighted by frequency of interactions) divided by the number of species in the network (69), with high connectance reflecting more generalized interactions in networks.

Trophic complementarity (TC) is a metric linked to function (host control) in antagonistic networks, with higher complementarity conferring higher host or pest control (34,74,75). TC defines the originality of each host morphotype relative to other host morphotypes based on their parasitoid assemblages. We calculated TC as the inverse of weighted NODF (nestedness) (TC) = (100 - NODF)/100 (as in 34).

Next, we calculated Nsal species-level metrics to uncover potential mechanisms of enemy loss under range expansions, including loss of Nsal specialist parasitoids, and weaker apparent competition by generalists. We estimated (Chao 1) richness of parasitoid morphospecies attacking Nsal (see S3). Next, we calculated the proportion of specialist parasitoids out of all parasitoids that attack Nsal (see above). Finally, as a weighted metric of specialization (analogous to (H_2_^’^)), we calculated d’ (species-level specialization) that represents how specialized parasitoid interactions with Nsal are given interaction frequencies of all parasitoid species in the network (73). We calculated species-level metrics for both generations separately and pooled (S1_Table S5).

To assess if parasitoids potentially fail to effectively switch from alternative hosts in the expanded range, we estimated apparent competition as Muller’s index (d_ij_) (76). This index (referred to as potential for apparent competition, PAC) calculates the likelihood that parasitoid (k) attacking host (i) developed in host (j), for all shared parasitoid species between host (i) and (j). d_ij_ summarizes the interactions between all paired hosts via all shared parasitoids and reflects PAC from host (j) to host (i), with 0 representing no shared parasitoids, and 1 high competition from host (j) to host (i). We treated Nsal as host (i) reflecting the strength of PAC from co-occurring hosts to the novel host, and calculated PAC as the sum of all hosts interacting with Nsal (for generations separately and pooled).

For each metric, we performed linear (LM) or generalized linear models (GLM) to compare metrics between regions at the site level. For linear models, we log-transformed some metrics to meet assumptions. For GLMs, we used Poisson, and negative binomial distributions in some instances to correct for overdispersion (S1_Tables S4,S5). Given that network size (species richness of hosts and parasitoids) correlates with network metrics and properties (77), we ran all analysis with and without network size (as an interaction term) to uncover if network size contributes to differences in network structure, or if mechanisms other than network size (i.e., changes in re-wiring of interactions) are influencing differences (77).

### Host-parasitoid interaction traits and phenology

We calculated peak parasitoid attack timing as the mean Julian date of when hosts were collected weighted by the number of emerged parasitoids from hosts collected on that date. We calculated mean Julian dates of parasitoid attack for each host morphotype collected at each site. Weighted mean parasitoid attack timing reflects when the host is vulnerable to parasitoids and the timing in which parasitoid morphospecies are attacking hosts. To compare parasitoid attack timing between Nsal and other hosts, we calculated effect sizes (as absolute values) as the log-response ratio (ln R) between each host morphotype and each generation of Nsal at each site (78). High effect sizes reflect phenological divergence in peak parasitoid attack timing between Nsal and other gall morphotypes. For each generation separately, we calculated average effect sizes of Nsal interactions with each gall morphotype at each site. We then calculated mean effect sizes of sites within regions ± 95% confidence intervals (C.I.). We note if C.I.s’ overlap 0 or not, reflecting phenological matching or divergence respectively between the host community and the range-expanding host. We ran LMs to compare effect sizes between regions for each focal host generation.

We measured morphological traits that are related to the ability of parasitoids to attack hosts and hosts to defend parasitoids. For each parasitoid morphospecies, we measured 1-3 individuals per region per host morphotype. We measured body size from the tip of the thorax to the end of the abdomen, the area of the wing, length of external ovipositor, width of thorax, and size of tibia (all in mm) (45). Since body size correlates with all other traits, we divided trait measurements by body size. Some parasitoids have internal ovipositors (67) that we were unable to measure. Thus, we also performed the trait analysis (see below) without ovipositors included, and found no difference in our results (see Supplementary Methods S1, for details). We also measured gall morphotype traits important in defense, such as gall size, internal traits (e.g., woody, fleshy or hollow), and external traits (e.g., nectar-producing, wooly, textured). We scored gall traits using our own observations and other resources (see S1 for details) (53,60,66).

We performed a Principal coordinates analysis (PCoA) on the full (metanetwork) parasitoid community and on the full host community using Gower’s dissimilarity that is useful for a mix of continuous and binary or categorical variables (79). Then, we calculated functional (or “interaction”) trait spaces by projecting host morphotypes onto parasitoid morphospecies trait space (as in 80,81). For each study site, we calculated weighted interaction centroids as the weighted (by frequency of interaction) mean position of assemblages of parasitoid morphospecies that a host morphotype interacts with in parasitoid trait space for each host morphotype. We calculated the distance of each host morphotype in interaction trait space to each generation of the focal species for each site network (80,81). We calculated mean morphological distance of each host morphotype with Nsal (each generation separately) at each site and then the mean (± 95% C.I.) of sites for each region. We compared morphological divergence between regions using a LM, with higher averages representing higher morphological divergence in parasitoid assemblages interacting with other hosts compared to the focal host. Also, to examine which parasitoid traits influence interactions with hosts, and which host traits influence interactions with parasitoids, we plotted PCoA biplots reflecting parasitoid trait space, host trait space, and interaction trait spaces, along with traits of gall morphotypes and parasitoid morphospecies (see S1 for details and S2 Fig. S2, S3 for biplots, as in 53). Analyses were performed using the following packages vegan in R (71).

### Linking network structure, function and traits

To examine how variation in network structure, function and traits are related, we performed a correlation analysis among network metrics relating to network structure and function, Nsal-specific metrics relating to mechanisms of enemy loss, and morphological and phenological trait divergence. Since TC is linked to function in antagonistic networks (34), we highlighted correlations between other metrics with this metric. To constrain the number of factors in this analysis, we used Nsal combined metrics (gamic and agamic generation combined) (see S1_Tables S4,5). We standardized all factors by calculating a z-score, and then performed a correlation analysis using corrplot in R (82), reporting which interactions were significant (*P* < 0.05).

### Biotic resistance: Nsal parasitoid attack rates

Finally, representing the function or the ability of host-parasitoids to provide biotic resistance to the novel host, we calculated parasitoid attack (or emergence) rates of Nsal. From our collections, we calculated parasitoid attack rates as the number of emergence holes from agamic galls (see S1_Supplementary Methods for details). To calculate parasitoid emergence rates more accurately, in 2021, we returned to sites in this study (along with three additional sites) and collected 500 Nsal (agamic) galls over their development (100 galls over 5 sampling periods). Each gall wasp placed in an individual gel capsule and kept in environmental chambers. We combined this data with collections we made using the same approach in 2007 and 2008 (18) (see S1). We performed a LM comparing parasitoid attack rates between the native and expanded range, including year as an interaction term.

## Results

### Host-parasitoid interaction networks

In total we reared 17,494 individual parasitoid wasps from collected gall wasp hosts, with 63 parasitoid morphospecies, and 14 host morphotypes (12 in which parasitoids were reared out of). At the network level, host-parasitoid networks were larger in the native range than in the expanded range (Chao 1: *P* << 0.001; Fig. 1); see full statistical results in S1_Table S4, with more host morphotypes (Chao 1: *P* = 0.035; Fig. 2a) and more parasitoid morphospecies (*P* = 0.025; Fig. 2b) (observed richness for host morphotypes and parasitoid morphospecies showed similar results, see S3). However, there was no difference in the number of interactions (Chao 1: *P* = 0.956), Shannon’s diversity (*P* = 0.781), or interaction evenness (*P* = 0.186) between regions.

**Figure 2:**
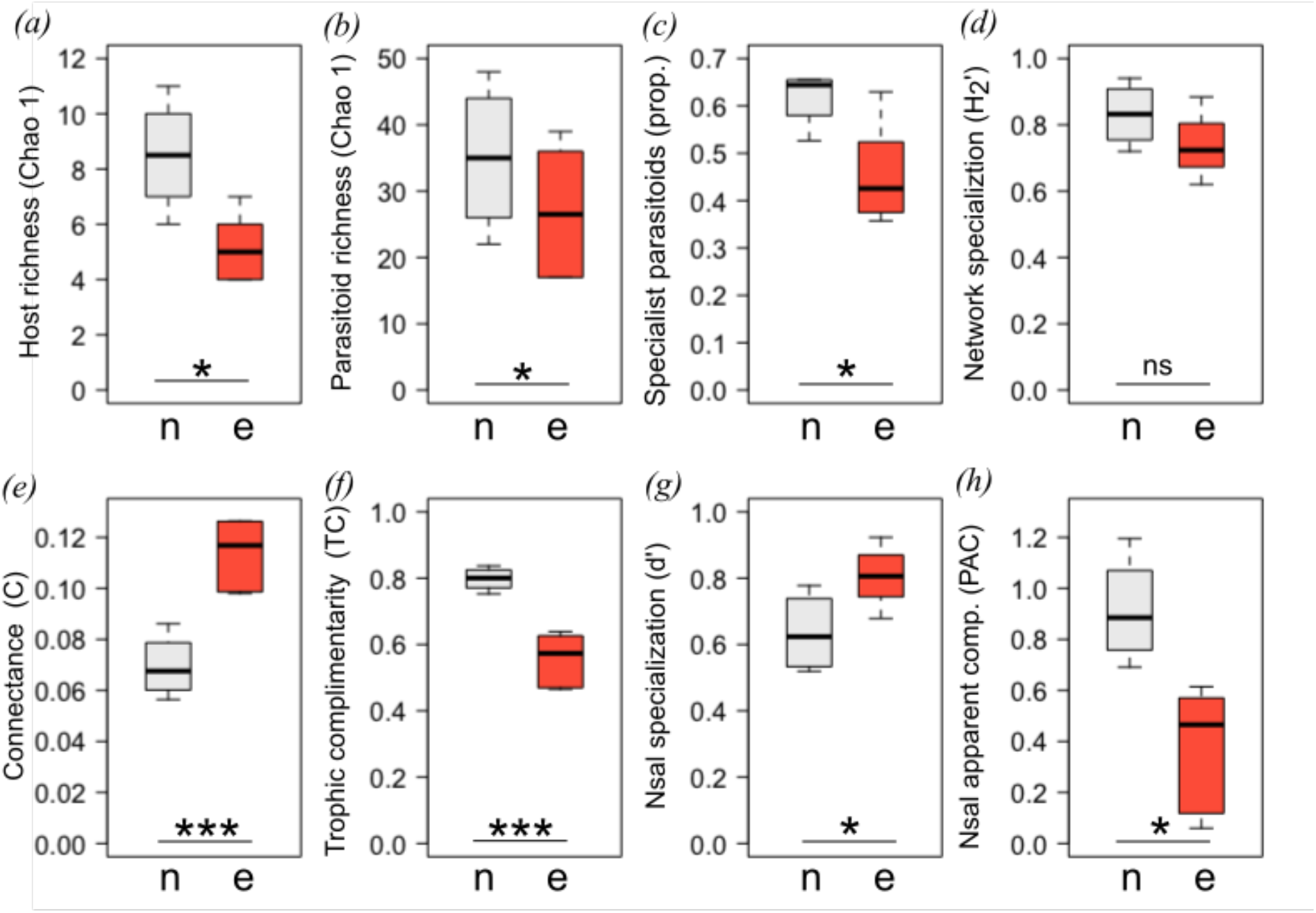
Host-parasitoid (a-f) network-level metrics and (g,h) species (Nsal)-level metrics. a) Estimated (Chao 1) number of host species, b) estimated (Chao 1) number of parasitoid species, c) proportion of specialist parasitoid species, d) network specialization (H2’), e) weighted connectance, f) trophic complementarity (TC), g) Nsal specialization (d’), h) potential for apparent competition (PAC) with Nsal in the native (n=4, gray) and expanded range (n=6, orange). Box plots depict the median +5^th^, 10^th^ and 25^th^ percentiles. Statistics are shown in Table S4,5 in S2. \**P* < 0.05,\*\**P* < 0.001,\*\*\**P* < 0.001.

Interaction networks had a higher proportion of specialist parasitoids attacking hosts in the native range (*P* = 0.028; Fig. 2c), which was also higher when including network size in the model (*P* = 0.040) (S1_Table S4). However, there was no difference in network specialization (H_2_’) between regions (*P* = 0.169) (Fig. 2d). Weighted connectance was higher in the expanded range (*P* < 0.001; Fig. 2e), with more shared partners for both host morphotypes (*P* = 0.001) and parasitoid morphospecies (*P* = 0.007) (S1_Table S4). Again, these metrics were still different when accounting for network size. Trophic complementary was higher in the native range (*P* < 0.001), including when accounting for network size, suggesting less overlap in parasitoid assemblages among host morphotypes in the native range (Fig. 2f).

For the Nsal species-specific metrics, Nsal had higher specialization (d’) in the expanded range compared to the native range (*P* = 0.007; Fig. 2g), which could be influenced by higher Nsal parasitoid richness (Chao 1 *P* = 0.002, S3). Collections of Nsal was much higher in the expanded range where this species is outbreaking, suggesting that estimates of Nsal parasitoid richness might be inflated by sampling intensity (see S3). The potential for apparent competition (PAC) was lower in the expanded range for hosts competing with Nsal through shared parasitoids (*P* = 0.007, Fig. 2h).

### Host-parasitoid interaction traits and phenology

Mean effect sizes of parasitoid attack timing (phenological divergence) was higher in the expanded range compared to the native range for the early spring gamic generation of Nsal(g) (*P* = 0.029; Fig. 3a; S1_Table S5), suggesting that peak parasitoid attack timing was more different between other hosts and the Nsal gamic generation in the expanded range compared to the native range. There was no difference in effect sizes of parasitoid attack timing between other hosts and Nsal(a) between regions (*P* = 0.654; Fig. 3b). Nsal(g) and Nsal(a) are on average further from other hosts in interaction trait space (morphological divergence) in the native range compared to the expanded range, with also more variation in the native range (gamic: *P* <0.001; agamic: *P* = 0.0003; Fig. 3c,d). Hosts are attacked by parasitoids with different body sizes, with more overlap in small and medium parasitoids attacking shared hosts (S2_Fig. S2). Parasitoids attack hosts of different gall size, and with different internal gall tissue (woody, hollow or fleshy), with external traits seeming to not strongly influence parasitoid assemblages (S2_Fig. S3).

**Figure 3:**
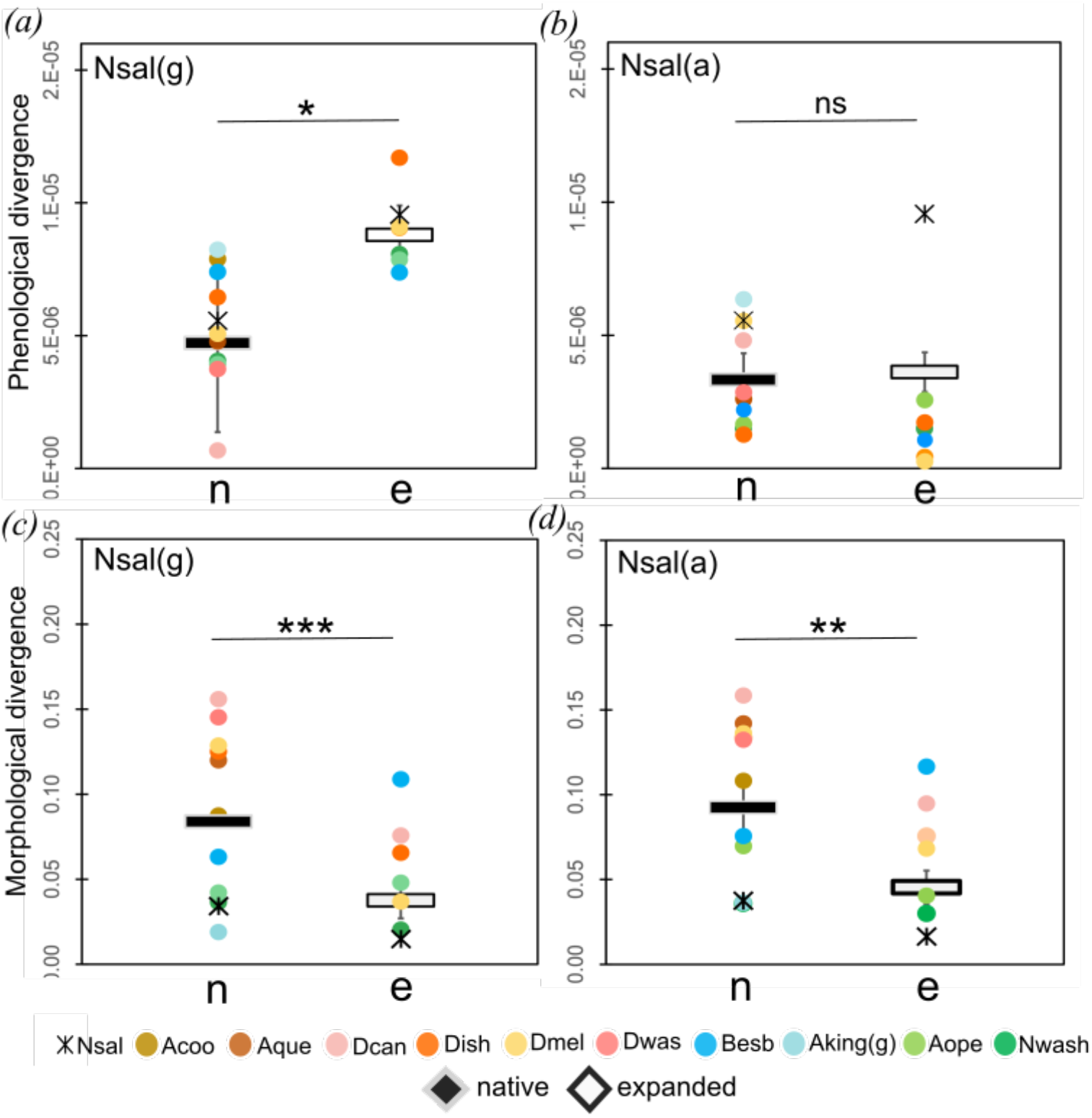
Host and parasitoid phenology (a,b) and morphology (b,c). (a,b) Effect sizes (ln(R)) of peak parasitoid attack timing for each host species relative to (a) Nsal(g) and (b) Nsal(a) at each study site in the native and expanded range. Mean (± 95% C.I.) of average attack times of sites in the native range (dark symbols) and expanded range (light symbols) are shown. When C.I.’s overlap zero, there is higher phenological matching in parasitoid attack timing of gall wasp hosts relative to the focal host. **(c**,**d)** Differences between centroids of each host species to focal species(a) Nsal(g) and (b) Nsal(a) of interacting parasitoids in parasitoid morphological trait space at each study site. Mean (± 95% C.I.) of average differences of sites in the native range (dark symbols) and expanded range (light symbols) are shown. Lower means represent greater overlap between other hosts and the focal host in interacting parasitoid morphological trait space (higher trait matching). Host species are depicted by different colors.

### Linking network structure, function and traits

Trophic complementarity (TC) had strong negative correlations with connectance (*R* = -0.95, *P* < 0.001; Fig. 4; S1_Table S6), both representing less specialized or more generalized interactions (and interaction overlap) in the expanded range. There was no relationship between specialization of Nsal (d’) and TC (*R = -0*.*38, P = 0*.*279*), but there was a positive trend between potential for apparent competition (PAC) and TC (*R* = 0.59, *P* = 0.07). Morphological divergence had a strong negative correlation with TC, showing the opposite of what we predicted that trait matching between hosts and parasitoids is related to decreased function (*R* = 0.92, *P* < 0.001). There was a negative trend between phenological divergence and TC (*R* = 0.53, *P* = 0.16), influenced by greater divergence of the gamic population with co-occurring hosts in the expanded range.

**Figure 4:**
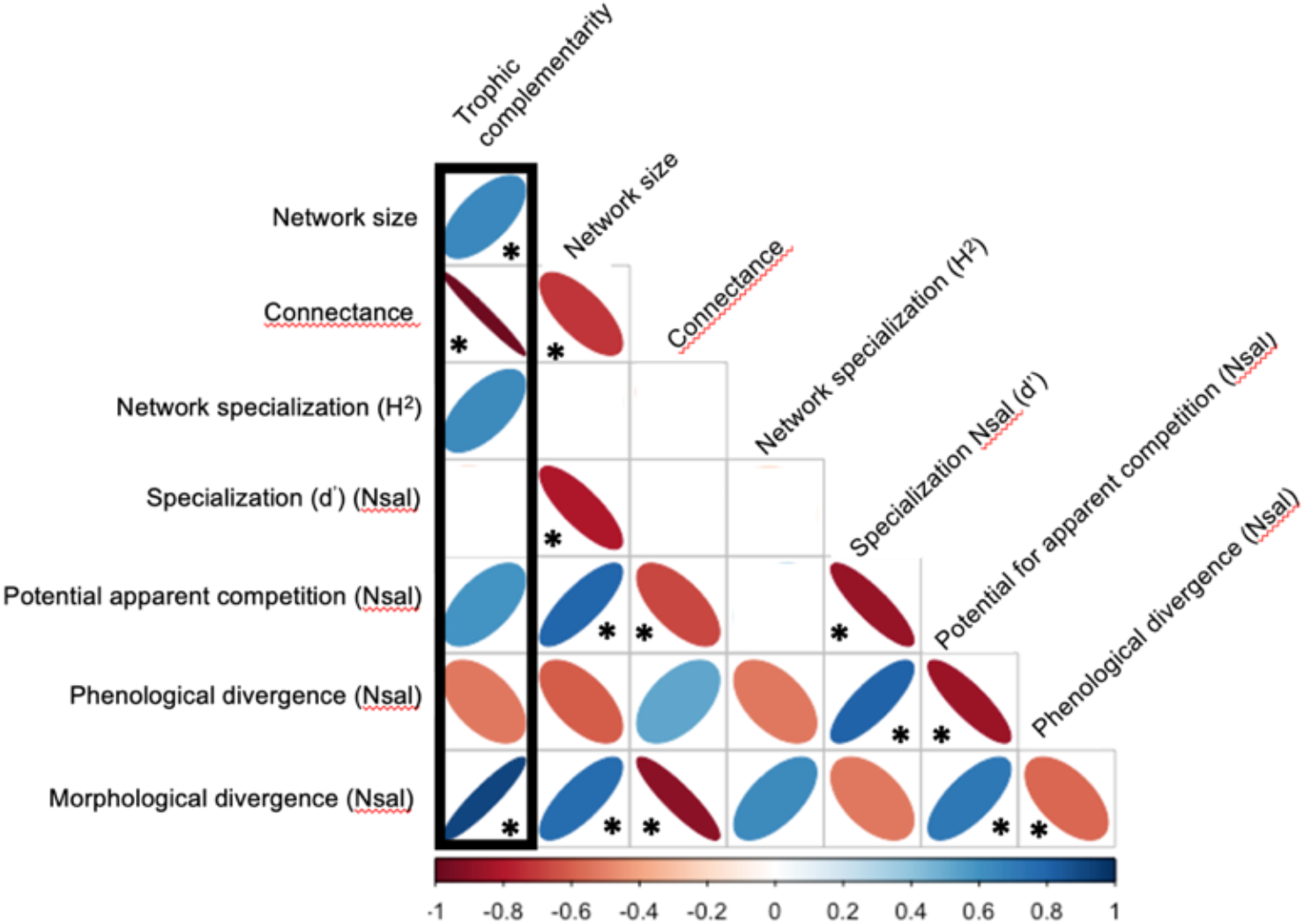
Relationships among host-parasitoid trait divergence and network structure. Shown are correlations (*R* > 0.50) among standardized factors. Blue ellipses show positive relationships, red negative relationships with the width and shade of ellipses reflecting strength of relationships (* *P* < 0.05) (S1_Table S6).

### Biotic resistance: Nsal parasitoid attack rates

We found no difference in the proportion of collected agamic galls that had emergence holes between the native and expanded range, although there is a trend for lower emergence holes in the expanded range (*P* < 0.329; Fig. 5a). Parasitoid attack rates (parasitoid emergence rates) were higher in the native range (*P* << 0.001), with no effect of year (Fig. 5b). We found similar results when only including the study sites that we created networks from (*P* < 0.001), but in this case the interaction with year was significant (range*year *P* = 0.01).

**Figure 5:**
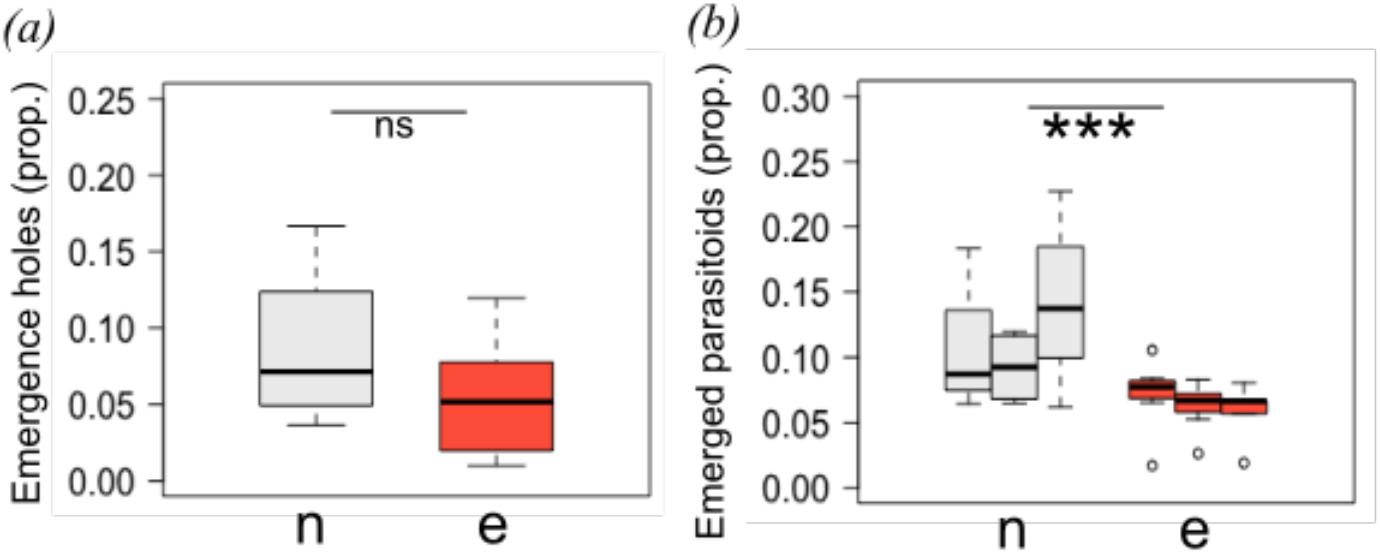
Nsal(a) parasitoid attack rates. a) Proportion of collected Nsal(a) galls with parasitoid emergence holes. b) Proportion of emerged parasitoids in Nsal(a) galls collected in 2007, 2008, 2021 (three bars) in the native (gray) and expanded range (orange). Box plots depict the median +5^th^, 10^th^ and 25^th^ percentiles. Statistics are shown in S5 \**P*<0.05,\*\**P*<0.001,\*\*\**P*<0.001.

## Discussion

Recipient oak gall wasp-parasitoid communities in the expanded range were less diverse, with fewer host morphotypes and parasitoid morphospecies. Whole networks in the expanded range were more connected, with fewer specialist parasitoids and with co-occurring hosts having less complementarity or turnover in parasitoid assemblages. Diverse, specialized host-parasitoid networks with higher complementarity are predicted to have higher function or host control (34,83). Interestingly, despite whole networks being more generalized in the expanded range, interactions between co-occurring hosts and Nsal were more specialized. Greater specialization of parasitoid assemblages on Nsal could result from Nsal hosting more specialist parasitoids or from lower potential for apparent competition (PAC) between co-occurring hosts and Nsal. That is, putative generalist parasitoids that attack multiple hosts may be more specialized (i.e., have unequal attack rates) on Nsal and co-occurring hosts in the expanded range. Our results suggest that differences in network structure of the recipient community and altered interactions with the novel host by putative generalist parasitoids may contribute to limited biotic resistance. These findings support that oak gall wasp-parasitoid communities are composed mainly of putative generalist parasitoids with broad host ranges that specialize (i.e., have high attack rates) on hosts with different morphological or spatio-temporal niches (24,42,63,84). Morphological divergence of parasitoid assemblages attacking co-occurring hosts and the novel host was not greater in the expanded range, reflecting that generalist parasitoids with similar traits attacked Nsal and co-occurring hosts. Phenological divergence was greater, suggesting that this could be a mechanism contributing to weaker apparent competition in the expanded range.

Poleward range-expanding species may experience weaker biotic interactions when they move into low diversity communities at the poles (21,23) except see (85). We found fewer host and parasitoid species in the poles and expanded range. Sites are smaller and patchier at the edge of the ecosystem’s range, and limited recruitment after the last glacial maximum could be one mechanism by which diversity decreases (23,86). Despite lower diversity in both groups towards the poles and the expanded range, there was a similar number of interactions. Networks were more connected with more overlap in parasitoid assemblages attacking co-occurring hosts. Higher network specialization of parasitoid assemblages in the native range could be driven by higher host diversity, with parasitoids specializing on hosts with differences in morphology, spatio-temporal niches, host immunity, or evolutionary divergence (42). While the expanded range is on an island, *Q. garryana-*ecosystems become naturally patchy at higher latitudes also on the mainland, with unsuitable habitat acting as a significant barrier between oak patches. Latitudinal patterns in diversity in *Q. garryana*-oak gall wasp and parasitoid communities follow similar trends on the mainland and when extended to the Island (23,87).

Lower trophic complementarity of parasitoid assemblages on hosts decreases host function (34,83). While low trophic complementarity (redundancy in parasitoid assemblages among hosts) is predicted to promote network stability or low variation in function, it is predicted to result in lower overall function or host control (34–36). Previous studies of antagonistic networks have found higher trophic complementarity leads to greater host control in networks (34–36), except see (75). Our results show that communities in the expanded range with lower trophic complementarity have lower host control (parasitoid attack rates) of the novel host, Nsal.

In addition to Nsal moving into less diverse and more generalized recipient communities in northern locations, range-expanding species often lose interactions when enemies (including specialist parasitoids) from the native range fail to shift (21,88). Opposite to this prediction, we found more putative specialist parasitoid species attacking Nsal in the expanded range. However, this could be a result from higher collections of Nsal in the expanded range, where it is outbreaking. *Amphidicous shickae* (Pteromalidae) is the most abundant specialist of Nsal that was initially described from Nsal and has not been recorded in any other samples of oak gall wasps (18,48,49,87), including in this study. This study and other findings suggest that *A. schickae* followed Nsal when it expanded its range to BC. In an earlier study, *A. schickae* attack rates were similar between regions (18). Here, we found higher attack rates in the expanded range, suggesting this species is equally effective at attacking Nsal in both regions (89). Thus, a loss of specialist parasitoids from the native range might not be a mechanism leading to weak biotic interactions in the expanded range.

Even though whole networks were more generalized in the expanded range, interactions with Nsal were more specialized. Greater specialized interactions (d’) with Nsal may be partially a result of higher Nsal attack rates by specialist parasitoids. However, generalist parasitoids are also more specialized in that they have greater asymmetrical frequency of attack between Nsal and other hosts in the expanded range. We found lower potential for apparent competition (PAC) between co-occurring hosts and Nsal in the expanded range, not due to fewer shared generalist parasitoid species but rather greater niche separation (unequal frequency of attack) between host species that shared parasitoids. This suggests that while generalist parasitoids can readily switch to attack Nsal, they may not do so effectively. Lower attack rates by generalist parasitoids could result from ineffective host switching or sharing between other hosts and Nsal when interactions are novel. Several mechanisms might lead to ineffective attack of novel hosts by locally adapted parasitoids, such as behavioral failure, physiological incompatibilities, or altered or novel parasitoid-parasitoid interactions (90–92). Other studies of range-expanding insects have found lower attack rates by generalist parasitoids where species have expanded their range (21,39).

One of the most abundant generalist parasitoids attacking Nsal, *Aprostoceus pattersonae* (Eupmelidae), had lower attack rates on Nsal in the expanded range (18). We do not know if generalist parasitoids attacking Nsal in the expanded range are native range populations that moved with Nsal or expanded range populations from other hosts (as some generalists were found in both regions). This information is critical to interpreting if lower attack rates by generalists result from ineffective switching by locally adapted populations or populations from the native range having lower efficacy in novel environments. These mechanisms of lower attack rates by putative generalists have occurred for introduced or range-expanding species (21,89). Uncovering pathways of parasitoid assembly on Nsal (as in 88) would be useful for future studies to uncover mechanisms of reduced generalist attack.

Our findings suggest that niche specialization by generalist parasitoids rather than loss of Nsal specialists might be important in determining variation in biotic resistance under range expansions. This finding supports that niche specialization by generalist parasitoids with broad host ranges is common in oak gall wasp-parasitoid communities, with richness in parasitoid communities maintained by partitioning of generalist parasitoids among different gall phenotypes (24,42,51). However, with increased molecular studies of parasitoid wasp communities and their interactions, more putative specialists originally described as generalists are being revealed (27,28,53,93,94). Identifying parasitoids via morphological features and taxonomic keys is challenging, and rearing out parasitoids from hosts may lead to incomplete information about associations. Future studies in this system will use molecular approaches to resolve interactions more accurately. Additionally, when creating networks, we likely miss interactions due to the window in which we made observations. We chose to sample the gall community when Nsal was developing on trees, but we could not capture associations for parasitoids emerging during other times in the season.

One mechanism of failed host sharing or switching may result from parasitoids in recipient communities lacking morphological adaptations to attack the novel host. We predicted that morphological trait divergence of parasitoids assemblages attacking other hosts and Nsal might be higher in the expanded range if trait mismatching is a mechanism influencing weak biotic resistance. However, we found that morphological divergence was lower in the expanded range, with traits of assemblages of parasitoids attacking co-occurring hosts and Nsal being similar. Given that networks were more connected and generalized in the expanded range, this result is not surprising. The native range contains more stem gall species that are large with tough exteriors. Parasitoids attacking these species have different traits, and their assemblages have little overlap with Nsal. Oak gall morphotypes that shared parasitoids with Nsal included fleshy leaf gall formers, *N. washingtonensis*, and *A. opertus* which are present in both regions, and small detachable species, such as *A. kingi*, that are not (23). Small generalists in the families Pteromalidae and Eulophidae are common in *N. saltatorius, N. washingtonensis*, and *A. opertus*.

Phenological divergence of assemblages of parasitoids attacking hosts (peak parasitoid attack timing) was higher in the expanded range for the earlier agamic generation. Greater phenological divergence of parasitoid attack timing in the expanded range was due to NSsal(g) being more apparent to parasitoids earlier than co-occurring species in the expanded range but not in the native range. Parasitoid attack timing is important for successful parasitism (47,91) and greater divergence between co-occurring hosts that share parasitoids could be a mechanism of low PAC in the expanded range.

Our snapshot natural experiment approach prevents us from comparing post to pre-invasion networks. As a result, we do not know if Nsal moved into less diverse, generalized networks or is creating less diverse, generalized networks. However, records of oak gall wasps on *Q. garryana* before the introduction of Nsal do not include many host species recorded in the native range (48,95), suggesting Nsal moved into a less diverse community.

We show that as species expand their range, they may move into structurally different networks and lose interactions with coevolved species or populations. While the number of interactions was resilient to network diversity changes, the distribution of interactions was not. Networks shifted from more specialized to generalized interactions between the lower latitude native range and higher latitude expanded range, which may result from lower species diversity and trait variation. Moving into less diverse, generalized networks might be typical for range-expanding species infiltrating recipient poleward communities. Additionally, interactions with range-expanding species may be lost, and we found that less effective interactions with putative generalist parasitoids might contribute to limited biotic resistance. Thus, variation in niche specialization by putative generalists, not interactions with specialists, might be important in creating high niche opportunities. This work provides novel insights into how population-level differences (local adaptation) might create open niches in short-distance-range expansions. Even when species move into very similar nearby habitats with similar species compositions, subtle differences in interaction networks may still have important consequences for population dynamics, potentially contributing to outbreaks and invasions.

## Supporting information

S3_Supplementary Methods_Rarefaction

## Acknowledgements

We thank Megan Blance and Kyle Skiver for help in the field, Catherine Ruis, Leslie Huang, Sarah Martin, Jesse Lofaso, Kelly McGourty, Julia Berliner, Tim Wong, Chris Johnston, Haley Hurst and Serena Feldman, Sabrina Bayshtock, and Gabriella Bayshtock helped in the lab. We thank landowners: OR Parks & Rec. Dept., OR Dept. of Fish & Wildlife, US Forest Service, The Nature Conservancy, City of West Linn Park, WA DNR, Weyerhaeuser, US Department of Defense, Canada Dept. National Defense, Saanich Parks, CRD Parks, BC Parks, and the Nature Conservancy Canada. Funding was provided by National Geographic Society (NGS-53395R-18) (KMP), National Science Foundation (DEB 1934387) (KMP), Binghamton University Clark Fellowship (DGJ), Binghamton University Provost Fellowship (AKM) and Binghamton University.

**Data and code will be made public upon publication**

