## Supplementary material for "Host-enemy interactions provide limited biotic resistance for a range-expanding species via reduced apparent competition": S1_Supplementary Methods and Tables

**S1_Supplementary Tables and Methods**

### Supplementary Methods

*Parasitoid enemy identification*

We first identified emergent wasps to family. Then for wasp families with known parasitoids, we identified individuals to the lowest taxonomic unit using taxonomic keys (1) and with the help of experts in some cases (2). We identified 63 unique parasitoid wasp morphospecies in 11 families in Superfamilies Chalcidoidea, Ichneumonoidea, and Platygastroidea (S1_Table S3). Inquiline wasps (i.e., cynipids that are not parasitoids, but feed on gall or plant tissue) also emerged from galls. Some inquiline wasps result in the death of the gall wasp, while others do not. Because we could not confirm how cynipid inquilines interacted with hosts, we did not include inquiline wasps in our networks. While we included morphospecies in families with known parasitoid wasps, not all wasps may have a direct interaction with the gall former host. For example, some species might be parasitoids of inquilines (wasp or non-wasp), or parasitoids of parasitoids (i.e., hyperparasitoids). While we removed non-wasp organisms interacting with galls (e.g., lepidopterans, coleopterans), it was impossible to remove all organisms from large and fleshy gall species (e.g., *Neuroterus washingtonensis, Andricus quercuscalifornicus*), and some parasitoids may have reared out these other hosts. In particular, wasps in the families Chalcididae and Ichneumonidae are described parasitoids of Lepidoptera (1). Given that many direct associations of parasitoids are undescribed, we took the conservative approach and included all morphospecies from families where wasps have been described as parasitoids, assuming that most caused deaths (be it direct or indirect) in gall wasp hosts. The vast majority of parasitoids reared were from taxonomic groups that are known to directly associate with gall wasps (1,3). For example, we only had one individual Chalcidoidea and 10 individual Ichneumonidae/Braconidae.

*Biotic resistance: Nsal parasitoid attack rates*

Finally, representing the function or the ability of host-parasitoids to provide biotic resistance to the novel host, we calculated parasitoid attack rates of Nsal. In our bulk collections, some parasitoids likely emerged before we collected galls. Thus, to calculate parasitoid attack rates we counted the number of emergence holes from agamic galls. We could distinguish parasitoid emergence holes in this generation from gall wasp emergences by the size of the hole (the gall wasp breaks open the side of the gall). Inquilines also create small emergence holes, but only two individuals emerged from our agamic Nsal collections. We could not estimate parasitoid attack rates using this approach for the spring gamic generation, because gall wasps and parasitoids create non-distinctive emergence holes. To calculate parasitoid emergence rates more accurately, in 2021, we returned to sites in this study (along with three additional sites) and collected 500 Nsal (agamic) galls over their development. We made collections of 100 galls 5X from early June to late July. We placed each gall into an individual gel capsule and kept capsules in environmental chambers. We combined this data with collections we made using the same approach in 2007 and 2008 (2). We performed a LM comparing parasitoid attack rates between the native and expanded range, including year as an interaction term.

*Morphological traits*

We measured morphological traits that are related to the ability of parasitoids to attack hosts and hosts to defend parasitoids. For each parasitoid morphospecies, we 1-3 individuals per region per host morphotype. We measured body size from the tip of the thorax to the end of the abdomen, the area of the wing, length of external ovipositor, width of thorax, and size of tibia (all in mm) (4). Since body size correlates with all other traits, we divided trait measurements by body size. Some parasitoids have internal ovipositors (1) that we were unable to measure and some of our morphospecies might have only reflected males. To account for this, we only included individuals with observable ovipositors for morphospecies where there was variation in ovipositor size (i.e., we excluded 0's). A handful of morphospecies has true 0's, with no replicate individuals with observable ovipositors. We also performed the trait analysis (see below) without ovipositors included, and found no difference in our results (see Table S4). We also measured gall morphotype traits that are likely important in defense (as in 5), we calculated gall volume ($cm^3$) and placed morphotypes in small, medium or large gall size categories. We scored the presence of internal defensive traits (if galls are woody, fleshy, hollow, or have radiating fibers) and external defensive traits (if galls are textured, wooly, secret nectar, possess spines, or come from a bract). We used our own observations and several resources to score traits (6,7).

We performed a Principal coordinates analysis (PCoA) on the full (metanetwork) parasitoid community and on the full host community using Gower's dissimilarity that is useful for a mix of continuous and binary or categorical variables (8). Then, we calculated functional (or "interaction") trait spaces by projecting host morphotypes onto parasitoid morphospecies trait space see (9,10). For each study site, we calculated weighted interaction centroids as the weighted (by frequency of interaction) mean position of assemblages of parasitoid morphospecies that a host morphotype interacts with in parasitoid trait space for each host morphotype. We calculated the distance of each host morphotype in interaction trait space to each generation of the focal species for each site network (9,10). We calculated mean morphological distance of each host morphotype with Nsal (each generation separately) at each site and then the mean ($\pm$ 95% C.I.) of sites for each region. We compared morphological divergence between regions using a LM, with higher averages representing higher morphological divergence in parasitoid assemblages interacting with other hosts compared to the focal host.

To uncover which parasitoid traits influence interactions with gall wasp morphotypes, we plotted parasitoid morphospecies in parasitoid space, and ("interaction trait space") or the non-weighted interaction centroids of interacting gall wasp morphotypes (from the meta-network) in parasitoid trait space. We color coded traits to reflect traits of parasitoids that influence interactions with gall wasps. We focused on body size (mm) (placing parasitoids in small, medium or large body size categories) as this trait is correlated with other parasitoid traits and had the strongest loading (S2_Fig. S2). Next to uncover what gall wasp traits influence interactions with parasitoid morphospecies, we plotted gall wasp morphotypes on gall trait space, and then the non-weighted interaction centroids of interacting parasitoids. We color coded traits representing gall size, internal traits, and external traits to visualize traits that influence parasitoids (S2_Fig. S3) (5).

Analyses were performed using the following packages vegan in R (11).

**Table S1:** Study site names and landowners in the native range (na) in Washington State (WA) and the expanded range (ex) Vancouver Island, British Columbia (BC).

| Site # | Site Name /Owner | Abbreviation | State /Province | Latitude/ Longitude | Site size (ha) | Habitat type |
|---|---|---|---|---|---|---|
| na1 | Fort Lewis, U.S. Department of Defense | FL | WA | 46.9170, -122.7084 | 129 | Woodland/ Grassland |
| na2 | Glacial Heritage Preserve, The Nature Conservancy | GHE | WA | 46.8646, -123.0366 | 78 | Grassland |
| na3 | Bald Hills, Weyerhauser | BAH | WA | 46.8098, -122.4359 | 9 | Grassland/ Rocky outcrop |
| na4 | Scatter Creek Wildlife Area, Washington Department of Natural Resources | SC | WA | 46.8331, -122.996 | 8 | Woodland/ grassland |
| ex1 | Little Mount Douglas, Saanich Parks | LMD | BC | 48.4892, -123.3539 | 32 | Rocky outcrop |
| ex2 | Mount Tolmie, Saanich Parks | MTO | BC | 48.4573, -123.3233 | 20. | Rocky outcrop/ woodland |
| ex3 | Rocky Point, National Defense Canada | RP | BC | 48.3249, -123.5408 | 29 | Woodland/ Rocky outcrop |
| ex4 | Cowichan Preserve, Nature Conservancy Canada | COW | BC | 48.8083, -123.6318 | 6 | Woodland |
| ex5 | Mt. Tzouhalem British Columbia Parks | MTZ | BC | 48.7896, -123.6386 | 10 | Rocky outcrop/ woodland |
| ex6 | Notch Hill, Nanoose Bay, National Defense Canada | NAN | BC | 49.2709, -124.1559 | 36 | Rocky outcrop/ woodland |

**Table S2:** Morphotypes (Hymenoptera: Cynipidae) collected from *Quercus garryana* trees at study sites, acronyms, regions and sites where they were found (see Table S1).

| **Cynipid morphotypes (species name), common name** | **Acronym** | **Gall type** | **Region(s)** | **Sites** |
|---|---|---|---|---|
| *Andricus coortus* Club gall wasp | Acoo | petiole, integral, monothalmous | Native, expanded | BAH, FL, GHE, SC, LMD |
| *Andricus kingi* (gamic) Red cone gall wasp | Akin(g) | leaf, detachable, monothalmous | Native | BAH, FL, GHE, SC |
| *Andricus kingi* (agamic) Red cone gall wasp | Akin(a) | leaf, detachable, monothalmous | Native | BAH, FL, SC |
| *Andricus opertus* Fimbriate gall wasp | Aope | leaf, integral, monothalmous | Native, expanded | All sites |
| *Andricus quercuscalifornicus* California gall wasp | Aque | stem, detachable, polythalamous | Native | BAH, FL, GHE, SC |
| *Besbicus mirabilis* Speckled gall wasp | Besb | leaf, detachable, monothalmous | Native, expanded | BAH, FL, GHE, SC, COW, LMD, MTZ |
| *Disholcaspis canescens* Round honeydew gall wasp | Dcan | stem, detachable, monothalmous | Native, expanded | BAH, FL, MTZ, NAN |
| *Disholcapis mamillana/ Disholcaspis simulata* Bullet gall wasp/Dried peach gall wasp | Dish | stem, detachable, monothalmous | Native, expanded | BAH, FL, GHE, SC, MTO, MTZ, NAN |
| *Disholcaspis mellifica* Twig gall wasp | Dmel | stem, detchable, monothalmousB | Native, expanded | BAH, FL, GHE, SC, COW, NAN, RP |
| *Andricus (Dros) pedicellatum* Hair stalk gall wasp | Aped | leaf, detachable, monothalamous | Native | FL |
| *Burnettweldia (Disholcaspis) washingtonensis* Round gall wasp | Dwas | stem, detachable, monothalamous | Native | BAH, FL, GHE, SC |
| *Neuroterus saltatorius* (gamic) Jumping gall wasp | Nsal(a) | leaf, integral, monothalamous | Native, expanded | All sites |
| *Neuroterus saltatorius* (agamic) Jumping gall wasp | Nsal(g) | leaf, detachable, monothalamous | Native, expanded | All sites |
| *Neuroterus washingtonensis* | Nwash | leaf, integral, polythalamous | Native, expanded | All sites |

**Table S3:** Parasitoid wasp morphospecies reared out of oak gall wasp hosts collected from study sites. Morphospecies were identified to the lowest taxonomic unit. Bolded morphospecies were reared out of Nsal galls and asterixis represent morphospecies only reared out of Nsal.

| Family | Morphospecies name | Identifier | Host morphotype | Native | Expanded |
|---|---|---|---|---|---|
| **Chalcidoidea** | | | | | |
| Chalcididae | *Conura* sp. | **P1** | Nwas | | X |
| Encyrtidae | *Habrolepis* sp. | **P2** | Dish | | X |
| Eulophidae | *Ephopalotus* sp. | **P3** | Nwas | | X |
| | *Chrysocharis* sp. | **P4** | Besb, Nwas | X | X |
| | *Euderus* sp. | **P5** | Nwas | X | X |
| | *Hemiptarsenus sp.* | **P6** | Aope | | X |
| | Eulophinae sp. 1 | **P7** | Nwas | | X |
| | *Pnigalio sp. | **P8** | **Nsal(g)** | | X |
| | ***Aprostocetus* sp. 1** | **P9** | Dcan, **Nsal(g), Nsal(a**), Nwas | X | X |
| | ***Aprostocetus* sp. 2** | **P10** | **Nsal(g),** Nwas | X | X |
| | ***Aprostocetus pattersonae*** | **P11** | **Nsal(g), Nsal(a**), Nwas | X | X |
| | ***Aprostocetus vericarri*** | **P12** | Aope, **Nsal(a), Nsal(g),**Nwas | X | X |
| | *Baryscapus* sp. 1 | **P13** | Dwas | X | |
| | *Baryscapus* sp. 2 | **P14** | Aque | X | |
| | Tetrastichinae sp. 1 | **P15** | Besb | X | |
| | **Tetrastichinae sp. 2** | **P16** | Aope, Dish, **Nsal(g), Nsal(a),** Nwas | X | X |
| Eupelmidae | ***Brasema* sp. 1** | **P17** | Aope, Acoo, Besb, Dish, Dmel, **Nsal(g), Nsal(a),** Nwas | X | X |
| | *Brasema* sp. 2 | **P18** | Dwas | X | |
| | *Eupelmus* sp. 1 | **P19** | Dish | X | |
| | *Eupelmus* sp. 2 | **P20** | Nwas | | X |
| | *Eupelmus* sp. 3 | **P21** | Aope | | X |
| Eurytomidae | ***Sycophila wilitzae*** | **P22** | Aope, Acoo, Besb, Dmel, **Nsal(g), Nsal(a),** Nwas | X | X |
| | *Sycophila* sp. 1 | **P23** | Acoo, Aque, Besb, Dwas Dcan, Dish, Dmel | X | X |
| | *Sycophila* sp. 2 | **P24** | Dmel | X | |
| | Eurytominae sp. 1 | **P25** | Nwash | X | X |
| | Eurytominae sp. 2 | **P26** | Dmel, Nwas | X | |
| | *Phylloxeroxenus* sp. | **P27** | Aope | X | |
| | *Eurytoma* sp. 1 | **P28** | Besb | X | |
| | *Eurytoma* sp. 2 | **P29** | Dcan, Dish, Dmel | X | |
| | *Eurytoma* sp. 3 | **P30** | Nwas | X | |
| | *Eurytoma* sp. 4 | **P31** | Dcan, Dmel | X | |
| | Eurytominae sp. 3 | **P32** | Dmel | X | |
| | Eurytominae sp. 4 | **P33** | Nwas | X | |
| | Eurytominae sp. 5 | **P34** | Nwas | | X |
| | Rileyinae sp. | **P35** | Nwas | | X |
| Ormyridae | ***Ormyrus sp.*** | **P36** | Aope, Dish, Dmel, **Nsal(a),** Nwas | | X |
| Pteromalidae | *Amphidocius schickae | **P37** | **Nsal(a)** | X | X |
| | ***Mesopolobus* sp.** | **P38** | Aope, Akin(g), **Nsal(g), Nsal(a),** Nwas | X | X |
| | Pteromalinae sp. 1 | **P39** | Nwas | X | X |
| | Pteromalinae sp. 2 | **P40** | Dmel | X | |
| | **Pteromalidae sp. 1** | **P41** | Aope, Aking, **Nsal(g), Nsal(a),** Nwas | X | X |
| | Pteromalidae sp. 2 | **P42** | Nwas | X | |
| | **Pteromalidae sp. 3** | **P43** | **Nsal(g),** Nwas | | X |
| | *Pteromalidae sp. 4 | **P44** | **Nsal(g)** | X | |
| | Pteromalidae sp. 5 | **P45** | Akin(g) | X | |
| | Pteromalidae sp. 6 | **P46** | Aope | | X |
| Torymidae | ***Bootanomyia dorsalis*** | **P47** | Aope, Besb, **Nsal(a**), Nwas | X | X |

| | | | | | |
|---|---|---|---|---|---|
| | *Torymus tubicola* | **P48** | Dwas | X | |
| | *Torymus* sp. 1 | **P49** | Aque, Nwas | X | |
| | *Torymus* sp. 2 | **P50** | Besb, Dish, Nwas | X | X |
| | *Torymus* sp. 3 | **P51** | Aque | X | |
| | *Torymus* sp. 4 | **P52** | Aque, Besb, Dish, Nwash | X | X |
| | *Torymus* sp. 5 | **P53** | Nwash | X | X |
| | *Torymus* sp. 6 | **P54** | Dwas, Aque | X | |
| **Ichneumonoidea** | | | | | |
| Braconidae | Braconidae sp. 1 | **P55** | Nwash | X | X |
| | Braconidae sp. 2 | **P56** | Nwash | | X |
| | Braconidae sp. 3 | **P57** | Nwash | X | X |
| Ichneumonidae | Ichneumonidae sp. 1 | **P58** | Nwash | | X |
| | Ichneumonidae sp. 1 | **P59** | Nwash | | X |
| | **Ichneumonidae sp. 1** | **P60** | Aope, **Nsal(g)** | | X |
| **Platygastroidea** | | | | | |
| Platygastridae | **Platygastridae sp. 1** | **P61** | **Nsal(a),** Nwash | X | X |

**Table S4:** Linear model (LM) and generalized linear model (GLM) output comparing network-level metrics between regions. Statistical results shown when models exclude and include network size. Analyses were performed with surveys combined (sites as replicates) and with surveys as replicates. F values (LM) or residual deviance (GLM) and p values (italics) shown. Bold represent significant differences, and (na) or (ex) which region has higher values.

| Metric | site network (region) | site network (region +richness) |
|---|---|---|
| Host richness (Chao 1) (GLM: Poisson) | **2.85, *0.045* (na)** | |
| Parasitoid richness (Chao 1) (GLM: Poisson) | **28.82, *0.025* (na)** | |
| Interactions (Chao 1) (GLM: Negative-binomial) | 10.11, *0.956* | |
| Network size (Chao 1) (GLM: Negative-binomial) | **10.22, <<0.001 (na)** | |
| Shannon Diversity (LM) | 0.08, 0.*781* | 0.08, *0.783* |
| Interaction Evenness (LM) | 2.09, *0.186* | 2.61, *0.145* |
| Specialist parasitoids (prop.) (LM) | **7.20, *0.028* (na)** | **6.30, *0.040* (na)** |
| H2' network specialization (LM) | 2.05, *0.189* | 1.82, *0.219* |
| Connectence (weighted) (LM) | **28.21*, <0.001* (ex)** | **29.99*, 0.001* (ex)** |
| Shared partners hosts (LM) | **27.86, *0.001* (ex)** | **25.13, *0.002* (ex)** |
| Shared partners parasitoids (LM: log-transformed) | **13.24*, 0.007* (ex)** | **11.60, *0.011* (ex)** |
| Trophic complementarity (LM) | **33.53, <*0.001* (na)** | **30.47, *0.001* (na)** |

**Table S5:** Linear model (LM) and generalized linear model (GLM) output comparing Nsal species-level metrics between regions. Statistical results shown when models exclude and include network size. Analyses were performed with surveys combined (sites as replicates) and with surveys as replicates. F values (LM) or residual deviance (GLM) and p values (italics) shown. Bold represent significant differences, and (na) or (ex) which region has higher values.

| Metric (Nsal) | site network (region) | site network (region +richness) |
|---|---|---|
| Parasitoid richness (Chao 1) (GLM: Poisson) | **2.85, *0.045* (ex)** | |
| Parasitoid specialists (prop.) (LM) | 3.02, *0.12* | 3.63, *0.098* |
| d' specialization (LM) | **6.43, *0.035* (ex)** | **9.01, *0.020* (ex)** |
| d' specialization (ga) (LM) | 1.97, *0.197* | 2.49, *0.158* |
| d' specialization (ag) (LM) | 1.07, *0.330* | 0.94, *0.365* |
| PAC (LM) | **13.05, *0.007* (na)** | **13.72, *0.006* (na)** |
| PAC (ga) (LM: log-transformed) | 4.62, *0.064* | 4.43, *0.07* |
| PAC (ag) (LM: log-transformed) | 1.10, *0.324* | 1.09, *0.331* |
| Phenological divergence (ga) (LM) | **7.06*, 0.029* (ex)** | |
| Phenological divergence (ag) (LM) | 0.216, *0.654* | |
| Phenological divergence (LM) | **6.43, *0.035* (ex)** | |
| Morphological divergence (ga) (LM) | **42.78, <*0.001* (na)** | |
| Morphological divergence (ag) (LM) | **35.11, <*0.001* (na)** | |
| Morphological divergence (LM) | **47.30, <*0.001* (na)** | |
| Morphological divergence (no ovipositor) (ga) (LM) | **35.14, *<0.001* (na)** | |
| Morphological divergence (ag) no ovipositor (LM) | **25.56*, <0.001* (na)** | |
| Morphological divergence no ovipositor (LM) | **37.01, *<0.001* (na)** | |
| Parasitoid emergence holes (ag) (LM) | **1.08, *0.329* (na)** | |

**Table S6:** Correlations between network-level, species-level, and trait metrics (R values and *p* values show). Bolded represent significant correlations

| | Trophic complementarity | Network size | Connectance | Network specialization (H') | Specialization Nsal (d') | PAC Nsal | Pheno-logical divergence (Nsal) |
|---|---|---|---|---|---|---|---|
| Network size | **0.64** ***0.046*** | | | | | | |
| Connectance | **-0.95** ***<0.001*** | **-0.71** ***0.021*** | | | | | |
| Network specialization (H2') | 0.63 *0.052* | 0.23 *0.522* | -0.47 *0.170* | | | | |
| Specialization Nsal (d') | -0.38 *0.279* | **-0.81** ***0.005*** | **-0.81** ***0.005*** | -0.15 *0.669* | | | |
| PAC Nsal | 0.59 *0.07* | **0.79** ***0.006*** | **0.79** ***0.006*** | 0.29 *0.410* | **-0.87** ***0.001*** | | |
| Phenological divergence (Nsal) | -0.53 *0.161* | -0.60 *0.064* | -0.60 *0.064* | -0.53 *0.118* | **0.80** ***0.005*** | **-0.85** ***0.002*** | |
| Morphological divergence (Nsal) | **0.92** ***0.0002*** | **0.77** ***0.009*** | **0.77** ***0.009*** | 0.63 *0.049* | -0.52 *0.119* | **0.72** ***0.019*** | **-0.58** ***0.081*** |
