## Supplementary material for "Host-enemy interactions provide limited biotic resistance for a range-expanding species via reduced apparent competition": S2_Supplementary Figures

**Supplementary Online Material**
**S2_Supplementary Figures**

**Host-enemy interactions provide limited biotic resistance for a range-expanding species via reduced apparent competition**

Kirsten M. Prior, Dylan G. Jones, Shannon A. Meadley-Dunphy, Susan Lee, Alyson K. Milks, Sage Daughton, Andrew A. Forbes, Thomas H. Q. Powell

**Figure S1:** Oak gall wasp host-parasitoid metanetwork, with surveys and sites combined. The bottom bars represent morphotypes of collected oak galls, with the width representing relative abundances. Orange bars represent the gamic and agamic generations of the focal species (Nsal). Links and top bars represent parasitoid morphospecies reared out of collected gall, with widths representing relative emergence rates. Dark blue bars represent hyperspecialists (emerged out of one host species), medium blue specialists (emerged out of 2-3 host species), and yellow generalists (emerged out of > 3 host species). See S1_Table S2 for host acronyms and Table S3 for parasitoid identifiers.

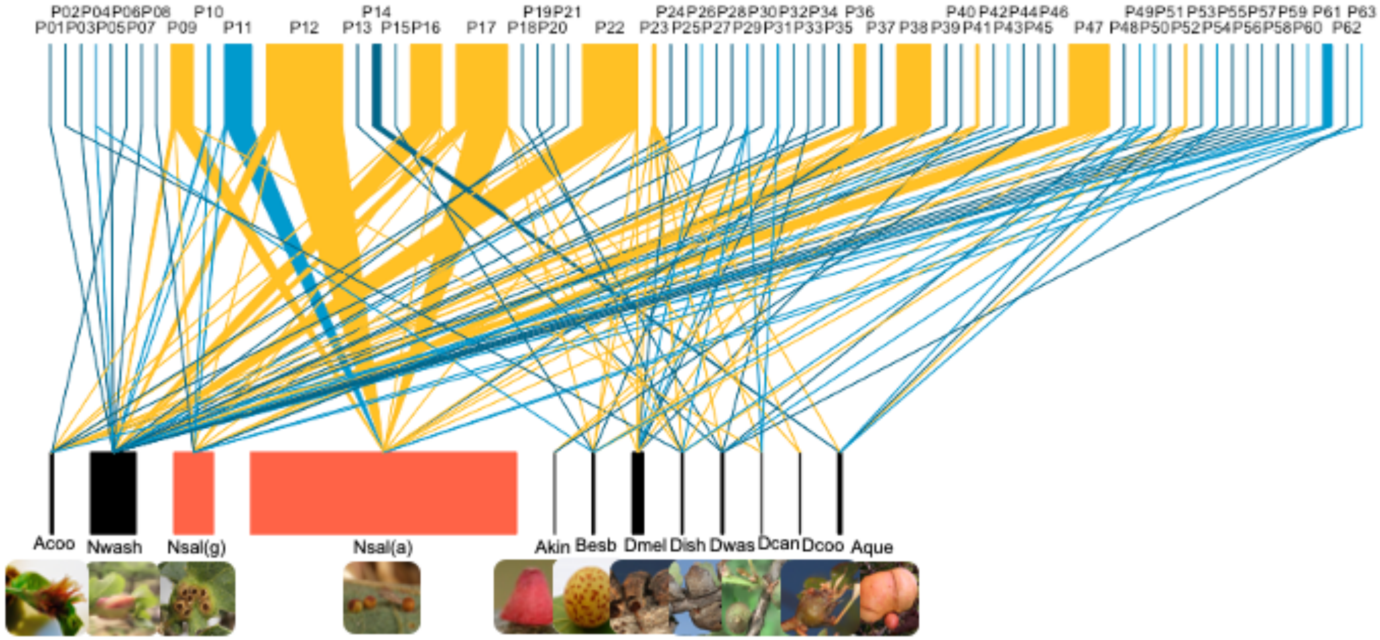

**Figure S2:** PCoA biplots of a) parasitoid morphospecies in parasitoid trait space. Colored symbols represent parasitoid morphospecies body sizes (mm) in three size categories; b) centroids of oak gall wasp morphotypes in parasitoid trait space (i.e., center of parasitoids galls interact with in parasitoid trait space). Colored symbols represent body sizes of interacting parasitoid morphospecies.

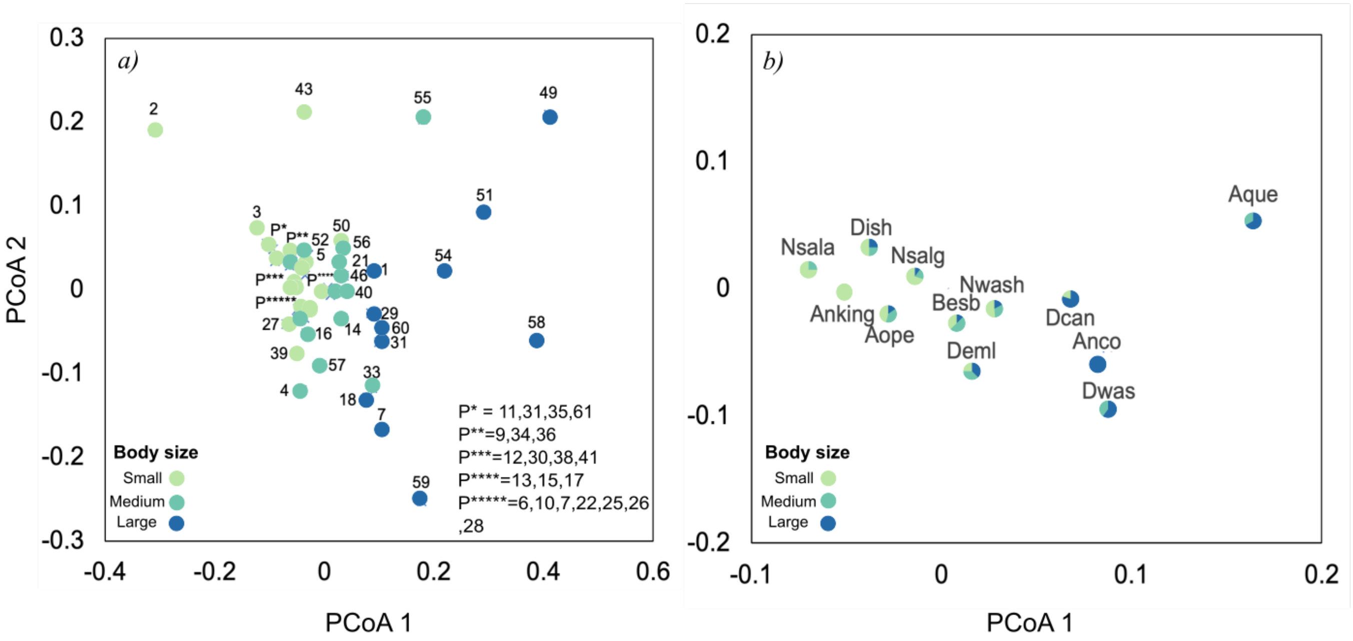

**Figure S3:** PCoA biplots of a-c) oak gall wasp morphotypes in in gall trait space. Colored symbols represent a) gall size, b) internal gall traits, and c) external gall traits; d-f) centroids of parasitoid morphospecies in gall wasp trait space (i.e., center of interacting parasitoids in gall trait space). Colored symbols represent a) gall size, b) internal gall traits, and c) external gall traits of interacting gall wasp morphotypes.

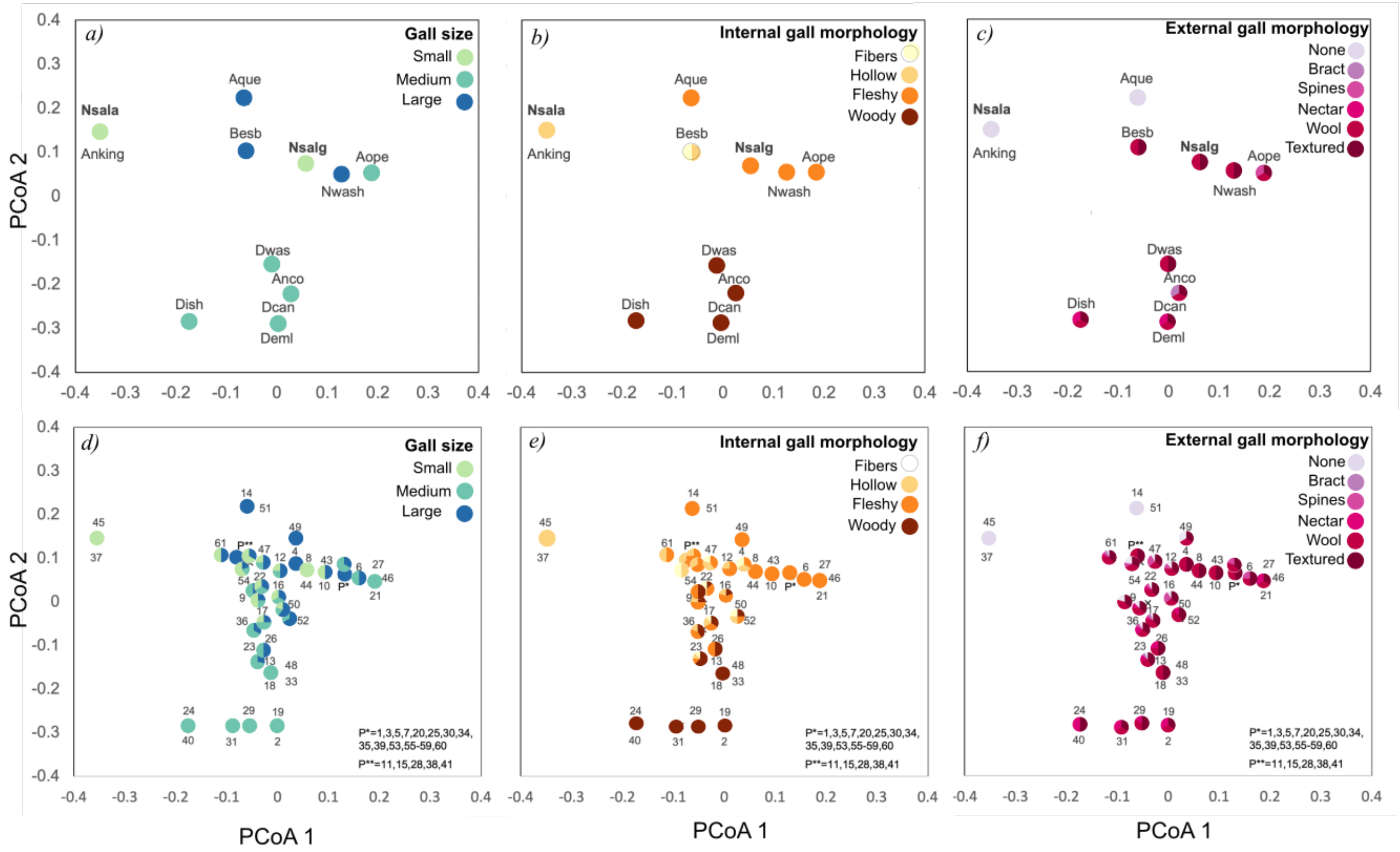
