## Supplementary material for "Host-enemy interactions provide limited biotic resistance for a range-expanding species via reduced apparent competition": S3_Supplementary Methods_Rarefaction

**Figure S5:** Regional rarefaction curves for the number of interactions and the number of parasitoid species.

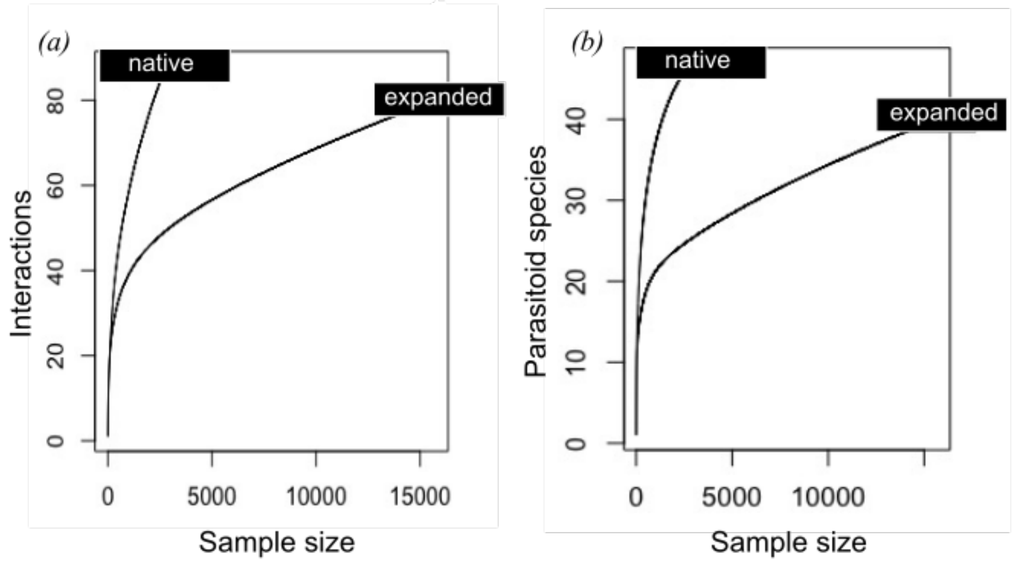


**Figure S6:** Mean (+/- SE) of observed (dark circles) and Chao 1 estimates (light triangles) of a) interaction richness, b) parasitoid richness, and c) cynipid morphotype richness at sites in the native and expanded regions.

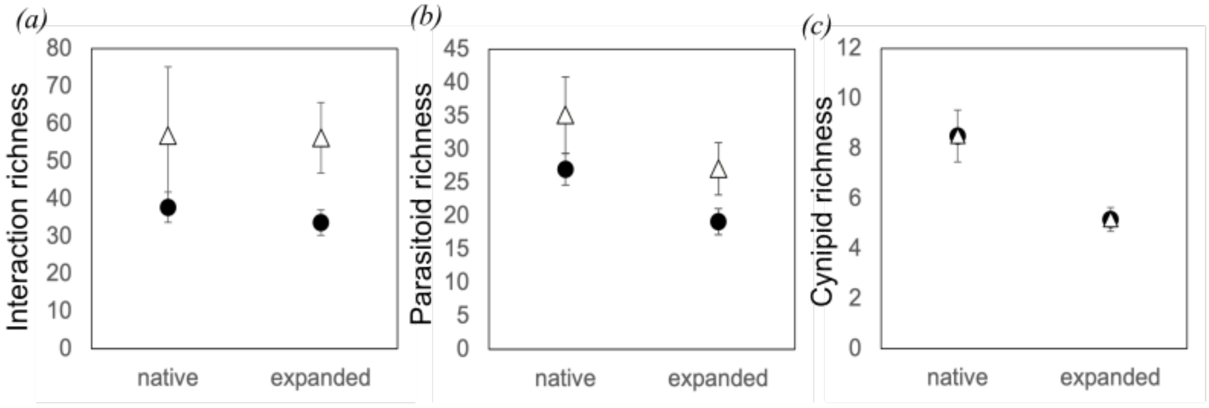

**Table S7:** Observed and Chao 1 estimates of interaction richness, parasitoid morphospecies richness, and cynipid morphotype richness at sites in the native and expanded range, along with means, and output of LM and GLM comparing observed and estimated richness between regions.

| Site | Interaction richness (Obs.) | Interaction richness (Chao 1) | Parasitoid richness (Obs.) | Parasitoid richness (Chao 1) | Cynipid richness (Obs.) | Cynipid richness (Chao 1) | Nsal parasitoid richness (Obs.) | Nsal parasitoid richness (Chao 1) |
|---|---|---|---|---|---|---|---|---|
| na1 | 48 | 111 | 20 | 22 | 11 | 11 | 6 | 6 |
| na2 | 40 | 43 | 29 | 30 | 9 | 9 | 4 | 4 |
| na3 | 33 | 42 | 29 | 40 | 6 | 6 | 4 | 4 |
| na4 | 30 | 32 | 30 | 48 | 8 | 8 | 7 | 7 |
| ex1 | 35 | 44 | 21 | 36 | 4 | 4 | 12 | 12 |
| ex2 | 29 | 36 | 16 | 17 | 5 | 5 | 9 | 9 |
| ex3 | 49 | 94 | 27 | 39 | 4 | 4 | 13 | 19 |
| ex4 | 26 | 37 | 14 | 17 | 5 | 5 | 9 | 9 |
| ex5 | 35 | 53 | 21 | 31 | 6 | 6 | 8 | 8 |
| ex6 | 28 | 73 | 16 | 22 | 7 | 7 | 7 | 8 |
| mean native | 37.7 | 56.9 | 27 | 35.1 | 8.5 | 8.5 | 5.2 | 5.2 |
| mean expanded | 33.7 | 56.3 | 19.2 | 27.1 | 5.2 | 5.2 | 9.7 | 10.8 |
| P-value | 0.96 | 0.39 | 0.011 | 0.025 | 0.035 | 0.045 | <0.002 | 0.045 |
| Res. Dev. | 10.11 | 9.75 | 8.43 | 28.82 | 2.85 | 2.85 | 8.42 | 2.85 |
